# Targetome analysis of malaria sporozoite transcription factor AP2-Sp reveals its role as a master regulator

**DOI:** 10.1101/2022.02.23.481739

**Authors:** Masao Yudaa, Izumi Kaneko, Yuho Murata, Shiroh Iwanaga, Tsubasa Nishi

## Abstract

Malaria transmission to humans begins with sporozoite infection of the liver. The elucidation of gene regulation during the sporozoite stage will promote the investigation of mechanisms of liver infection by this parasite and contribute to the development of strategies for preventing malaria transmission. AP2-Sp is a transcription factor (TF) essential for the formation of sporozoites or sporogony, which takes place in oocysts in the midgut of infected mosquitoes. To understand the role of this TF in the transcriptional regulatory system of this stage, we performed ChIP-seq analyses using whole mosquito midguts containing late oocysts as starting material and explored its genome-wide target genes. We identified 697 target genes, comprising those involved in distinct processes parasites experience during this stage, from sporogony to development into the liver-stage, and representing the majority of genes highly expressed in the sporozoite stage. These results suggest that AP2-Sp determines basal patterns of gene expression by targeting a broad range of genes directly. The ChIP-seq analyses also showed that AP2-Sp maintains its own expression by a transcriptional auto-activation mechanism (positive feedback loop) and induces all TFs reported to be transcribed at this stage, including AP2-Sp2, AP2-Sp3, and SLARP. The results showed that AP2-Sp exists at the top of the transcriptional cascade of this stage and triggers the formation of this stage as a master regulator.

## Introduction

Malaria is still a great threat to humans, killing nearly half a million people per year(1). Malaria parasites possess a complex lifecycle. The capacity of these parasites to infect different target organs and adapt to distinct host environments is supported by mechanisms to create gene expression repertoires unique to each parasite stage. Thus, to understand the molecular mechanisms underlying this capacity, it is important to elucidate the mechanisms of stage-specific gene regulation.

The APETALA2 (AP2) family is the sole family of gene-specific transcription factors (TFs) in this parasite(2) and members of this family play central roles in stage-specific transcriptional regulation in malaria parasites(3)(4)(5). Recent genome-wide studies revealed that AP2 family TFs create gene expression patterns for each stage by targeting hundreds of genes directly(6)(7)(8)(9). Therefore, identifying the master TF that governs specific stages and determining its target genes are important strategies for elucidating mechanisms of infection in this parasite.

Sporozoites are motile forms that play a central role in parasite transmission to the vertebrate host(10). They are generated in the oocysts formed within the mosquito midgut and then migrate to the salivary glands. When a mosquito bites, the sporozoites are released into the skin and are transported to the liver via the circulation. There, they enter the liver stage to transform into merozoites that are erythrocyte-invasive forms (11). Studies on gene regulation during this stage are important to understand the molecular mechanisms underlying parasite infection of the liver. Thus far, four TFs have been reported to play important roles in gene regulation during this stage: AP2-Sp, AP2-Sp2, AP2-Sp3, and sporozoite and liver stage asparagine-rich protein (SLARP)(5)(12)(3). However, studies on these TFs including their target genes and their interrelationships are lacking, and their roles in transcriptional regulation during the sporozoite stage remain to be elucidated. The likely reason for this paucity of research is the necessity to obtained sporozoites by dissecting infected mosquitoes(10). Genome-wide studies on these TFs, particularly ChIP-seq analyses to determine their binding sequences and target genes, have been hampered due to the small number of sporozoites obtained from infected mosquitoes. Thus, the question of how the gene expression repertoire unique to this stage occurs has been left unanswered.

AP2-Sp is a TF that is essential for formation of the sporozoite stage(5). The expression of AP2-Sp is first observed in the nuclei of oocysts approximately seven to ten days after an infective blood meal of a mosquito, i.e., a few days earlier than the onset of sporozoite formation or sporogony in the oocyst. AP2-Sp expression then continues through the sporozoite stage. Depletion of this gene arrested oocyst development before sporogony, and no sporozoites were produced in the oocyst. The AP2-domain of this TF binds the six-base motif [C/T]GCATG. Many genes important for sporozoite functions harbor this binding motif in their upstream regulatory regions, where this motif functions as a *cis*-acting element of the gene promoters. These findings have suggested an important role of this TF during the sporozoite stage. However, it remains unclear to what extent this TF contributes to gene regulation during this stage and how it interconnects with other concomitantly expressed TFs.

In this study, we aimed to demonstrate the role of AP2-Sp in *Plasmodium berghei* as a master regulator by investigating the whole range of targets or the “targetome” of this TF by ChIP-seq. To circumvent the problems of the low recoveries of sporozoites obtained from mosquitoes, we used whole mosquito midguts containing oocysts under sporogony as the starting materials for ChIP-seq. The results comprehensively demonstrate that AP2-Sp regulates genes expressed during this stage; its targets broadly encompass genes for sporogony, salivary gland infection, and hepatocyte infection. We also show that AP2-Sp directly induces other TFs expressed during this stage, playing the role of a stage-specific master gene regulator.

## Results

### ChIP-Seq of AP2-Sp using mosquito midguts containing oocysts

In ChIP-seq, the number of cells utilized in the assay is an important determinant of data quality because the maximum read depth achieved by ChIP from a single haploid cell is only one on a given genome locus. In previous ChIP-seq studies, we used five to ten hundred million haploid parasites to obtain nanogram quantities of DNA by ChIP. To attain this number in the sporozoite stage (in our laboratory, no more than fifty thousand oocyst sporozoites and ten to twenty thousand salivary gland sporozoites are obtained from one mosquito), it is necessary to dissect at least ten thousand infected mosquitoes. Thus, performing ChIP-seq analysis of AP2-Sp in this stage was unrealistic.

To solve this problem, we conceived using oocysts as the starting material. After an infective blood meal, parasites transform into round oocysts on the basal lamina of the midgut. The oocysts grow, replicating their genome, and after 10–14 days, divide into multinuclear cells, called sporoblasts, which act as centers of sporogony. Hundreds of sporozoites bud promptly from each sporoblast. AP2-Sp expression begins just before sporoblast formation and then is localized in the nuclei aligned beneath the plasma membrane of sporoblasts (5). If an average of 100 oocysts is formed in the mosquito midgut and each oocyst contains ten thousand nuclei, one midgut is estimated to contain nuclei equivalent to 1 × 10^6^ haploid parasites, and 100 mosquitos would satisfy the required number of nuclei described above. However, one drawback is that the oocysts cannot be purified from the mosquito midgut, and thus, to perform this experiment, whole midguts must be used as the starting material. This means that the samples will contain considerable amounts of contaminating proteins and genomic DNA derived from the mosquito host. We anticipate that the efficiency of ChIP will be diminished under such conditions, which will affect the data quality deleteriously.

Therefore, we performed the ChIP-seq experiments using infected midguts and assessed the data quality. In the first experiment, we harvested 450 *Anopheles stephensi* mosquito midguts for ChIP-seq at 14 days after an infective blood meal, when AP2-Sp expression was observed in the majority of oocysts. In the midguts, due to asynchronous oocyst development (13), there were oocysts at various developmental stages, including pre-sporogony oocysts, oocysts undergoing sporogony, and oocysts containing sporozoites. Based on the estimation described above, these oocysts were estimated to contain 4.5 × 10^8^ parasite nuclei.

The harvested midguts were immediately fixed with 1% paraformaldehyde and stored in the fixation solution until all mosquitoes were dissected. The fixed midguts were then subjected to ChIP with anti-GFP antibodies. Sequencing of the input DNA, which was extracted from the midgut-lysate, showed that ratio of the parasite genome to the *Anopheles stephensi* genome in the original sample was approximately 1: 3.1, indicating that the starting sample contained a substantial amount of genomic DNA from the parasites. By peak-calling using ChIP-seq data, 1,270 peaks were called (Table S1A, FDR < 0.01, fold enrichments > 3).

To investigate the binding sequences *in vivo*, regions within 100 bp from the peak summits were extracted, and the sequences around the summits were analyzed. The results demonstrated that the motif [T/C]GCA[T/C][G/A] (and its reverse complementary motif [T/C][G/A]TGC[G/A]) was enriched around the summits (Fig. 1A and Table S1B). This motif is identical to the one that we reported in our previous paper, [T/C]GCATG, except that the current motif contains TGCACA and TGCATA as minor variants that were not covered in our previous study using electrophoretic mobility shift assays(5). Of these peaks, 90.6% contained at least one motif sequence within their peak regions, and the average distance of the predicted summit to the nearest motif was 54.0 bp (Fig. 1B). The mapped view of reads showed that representative sporozoite-specific genes, including the AP2-Sp targets predicted in our previous paper (5), harbor clear AP2-Sp peaks in their upstream regions (examples are shown in Fig. 1C). These results (i.e., high proportion of parasite DNA in the infected midgut and low background in the obtained ChIP-seq data) suggest that the efficiencies of ChIP-seq from infected midguts were higher than we initially supposed.

**Figure 1.**
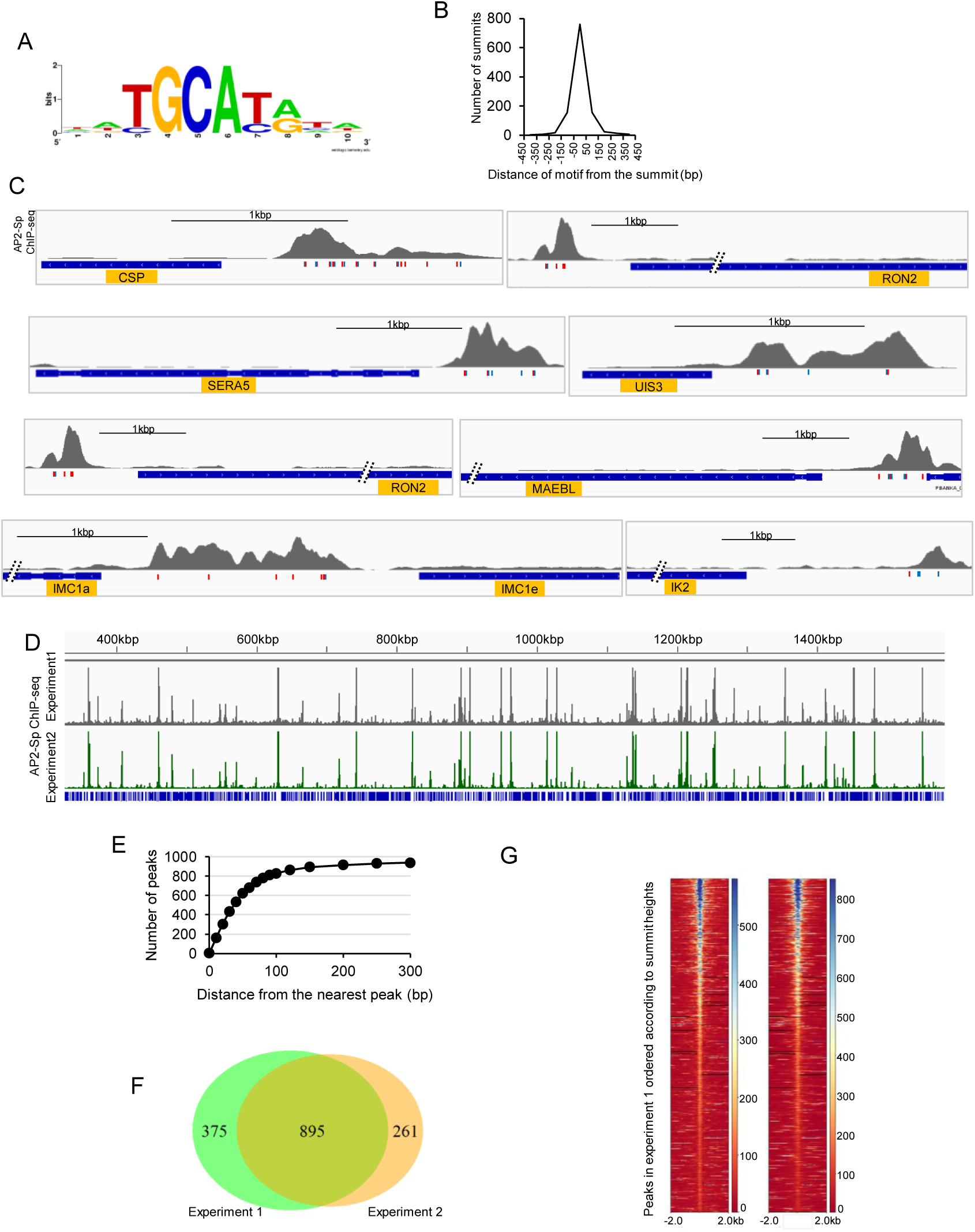
ChIP-seq of AP2-Sp. A. Motif associated with summits of AP2-Sp peaks in experiment 1. Binding sequences nearest to the summit were visualized with WebLogo software. B. Distance from the summit to the nearest motif sequence was calculated in each AP2-Sp peak, and the distribution is shown with a histogram. The horizontal axis indicates the distance from the summit to the nearest motif sequence (the bin size is 100 bp). C. ChIP-seq peaks of AP2-Sp upstream of target genes. Seven genes functionally different in the sporozoite stage were selected (see also Fig. 2D and Fig. 2E). Binding motifs of AP2-Sp under the peaks are indicated by bars. Peaks are visualized by Integrative Genomics Viewer (IGV) software. CSP, circumsporozoite protein; IMC1a, inner membrane complex protein 1a; SERA5, serine repeat antigen 5; MAEBL, membrane associated erythrocyte binding-like protein; SPECT2, sporozoite micronemal protein essential for cell traversal 2; RON2, rhoptry neck protein 2; UIS3, upregulated in sporozoite 3; IK2, eukaryotic translation initiation factor 2-alpha kinase 2 D. Comparison of the mapped views of ChIP-seq peaks obtained by two independent experiments. Reads were mapped on chromosome 14. E. Numbers of common peaks between experiment 1 (1270 peaks) and experiment 2 (1156 peaks) within the selected distance were plotted. The plot indicates that approximately 80% (895 peaks) of the peaks in experiment 2 had counterparts in experiment 1 within 150 bp. F. Venn diagram showing common peaks of two ChIP-seq experiments. Peaks whose summits were within 150 bp were regarded as common. G. Coverage maps of two ChIP-seq experiments were created by setting positions of peak summits identified in experiment 1 in the center.

Therefore, we next performed ChIP-seq using a smaller number of mosquitoes, i.e., 190 mosquitoes. In this experiment, the ratio of parasite DNA to *A. stephensi* DNA in the input DNA was 1:0.96. By peak call, 1,156 peaks were identified (Table S2A), and the motif around the peak summits was identical to that obtained with the former experiment (Table S2B). Of these peaks, 95.9% contained at least one motif sequence within the peak region, and the average distance of the predicted summit to the nearest motif was 53.3 bp. The comparison of graphical views of the ChIP-seq peaks between these experiments is shown in Fig. 1D. Between these experiments, there were 895 common peaks (peak summits were within 150 bp) (Fig. 1E, Fig. 1F, and Fig. G), demonstrating that the results were highly reproducible.

### Comprehensive activation of host cell invasion-related genes by AP2-Sp

To investigate the role of AP2-Sp in transcriptional regulation in the sporozoite stage, the genome-wide targets of AP2-Sp were predicted. The prediction assumed that the binding sites of a TF identified by ChIP-seq exist within 1,200 bp of the first methionine codon of the gene. This criterion produced useful results in our previous studies, and its validity is also supported by our previous observation that in many sporozoite-specific genes, the binding sequences of this TF were located within this region (5). Using this criterion, 784 and 850 genes were predicted as targets from the peaks obtained via experiments 1 and 2, respectively (Table S1C and Table S2C), and 697 genes were common between them (Table S3A and Fig. 2A). These common genes were used for the following analyses.

**Figure 2.**
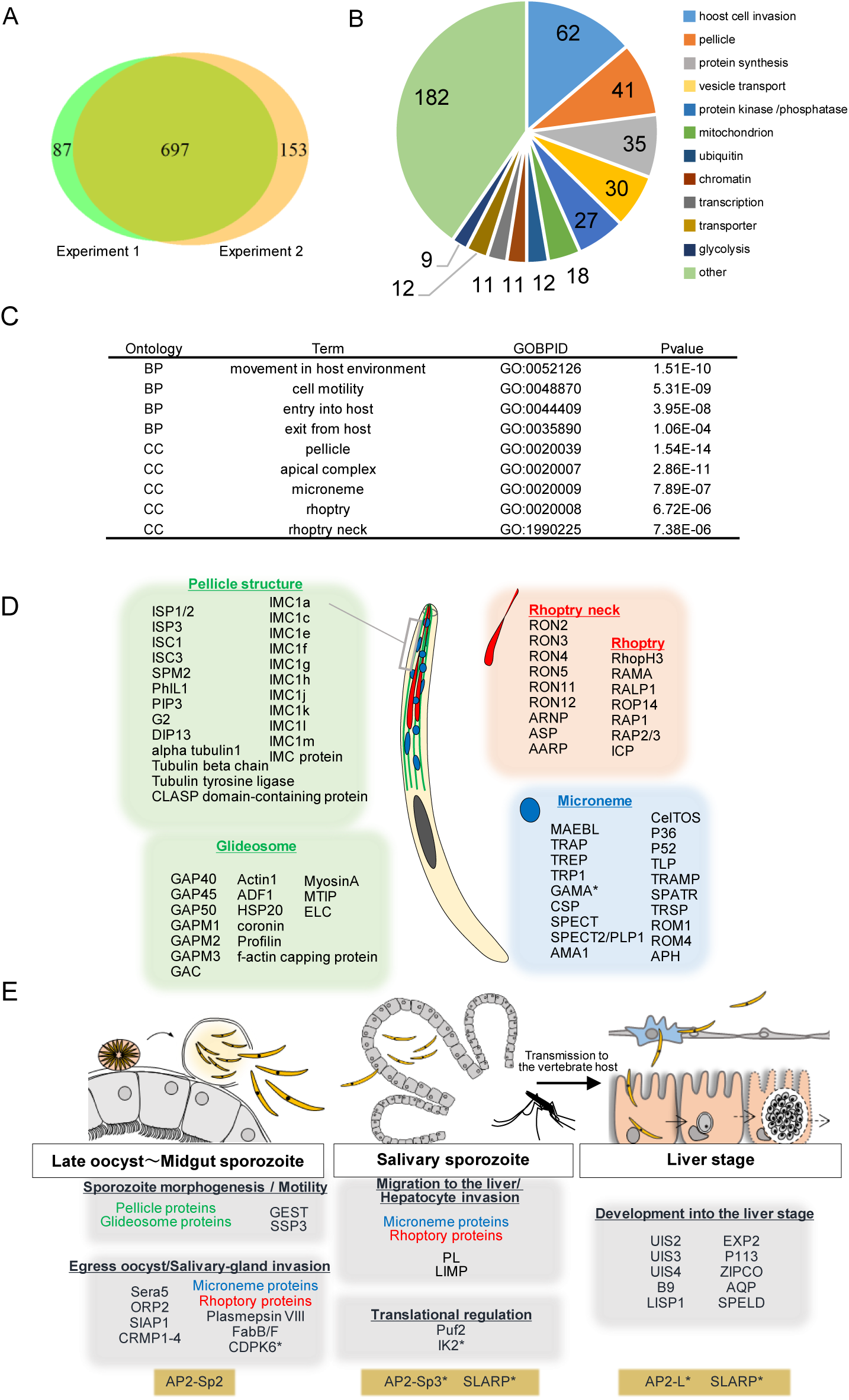
Targetome of AP2-Sp. A. Common target genes predicted by two ChIP-seq experiments B. Common target genes were classified according to functional categories. The number of target genes in each group is shown on a pie graph. C. GO enrichment analysis. Terms enriched in target genes (p<0.01) are listed. BP, Biological Process; CC, Cellular Component D. Products of target genes are grouped by localization to subcellular structures characteristic to motile stages: microneme, rhoptry, and pellicle structure. Proteins belonging to the pellicle structure are further classified into cytoskeletal proteins and proteins involved in gliding motility. The gen marked by an asterisk was additionally predicted as targets manually using ChIP-seq peaks and RNA-seq data. IMC, inner membrane complex; ISP, IMC sub-compartment protein; ISC, IMC suture component; PHIL1, photosensitized INA-labeled protein 1; SPM, subpellicular microtubules; GAP, glideosome-associated protein; GAPM, glideosome-associated protein with multiple-membrane spans; GAC, glideosome-associated connector; ADF, actin-depolymerizing factor;;MTIP, myosin A-tail interacting protein; ELC, myosin essential light chain: RON, rhoptry neck protein; ARNP, apical rhoptry neck protein; ASP, apical sushi protein; AARP, apical asparagine-rich protein; RhopH3, high molecular weight rhoptry protein 3; RAMA, rhoptry-associated membrane antigen; RALP1, rhoptry-associated leucine zipper-like protein 1; ROP14,rhoptry protein 14; RAP, rhoptry associated protein; ICP, inhibitor of cysteine proteases; MAEBL, merozoite apical erythrocyte-binding ligand; TRAP, thrombospondin-related anonymous protein; TREP, TRAP-related protein; TRP1, thrombospondin-related protein 1 GAMA, GPI-anchored micronemal antigen; CSP, circumsporozoite protein; SPECT, sporozoite microneme protein essential for cell traversal: PLP1, perforin-like protein1; AMA1, apical membrane antigen 1; CelTOS, cell transversal for ookinetes and sporozoites; TLP, TRAP-like protein: TRAMP, thrombospondin-related apical membrane protein; STAPR, secreted protein with altered thrombospondin repeat domain; TRSP, thrombospondin-related sporozoite protein; ROM, rhomboid protease; APH, acylated pleckstrin-homology domain-containing prote E. Parasite progression through the sporozoite stage, starting with sporogony and ending with development into the liver stage, is illustrated. Target genes are categorized according to the steps they are involved in. The four genes marked by an asterisk were additionally predicted as targets manually using ChIP-seq peaks and RNA-seq data. GEST, gamete egress and sporozoite traversal protein; SSP3, sporozoite surface protein 3; SERA, serine repeat antigen 5; ORP2, oocyst rupture protein 2; SIAP1, sporozoite invasion-associated protein 1; CRMP, cysteine repeat modular protein; PL, phospholipase; Puf2, Pumilio-FBF family protein 2; IK2, eukaryotic translation initiation factor 2-alpha kinase 2: UIS, upregulated in sporozoite; LISP1, liver specific protein 1; ZIPCO, ZIP domain-containing protein; AQP, aquaglyceroporin; SPELD, sporozoite surface protein essential for liver stage development

The classification of these targets according to functional categories is shown in Fig. 2B and Table S3B. Genes related to pellicle structure and host cell invasion were the most abundant in the predicted targets. Consistently, Gene Ontology (GO) term analysis revealed that terms such as “pellicle”, “cell motility”, and “apical complex” were significantly enriched (p<0.01) in the targets (Fig. 2C). Fig. 2D shows target genes related to these terms that are grouped based on the subcellular localization of the products: “pellicle cytoskeleton”, “glideosome”, “microneme”, “rhoptry”, and “rhoptry neck”. We also classified the target genes according to their involvement in the steps that parasites undergo during this stage: “sporogony”, “egress oocyst and salivary-gland invasion”, “migration to the liver and hepatocyte invasion”, and “development into the liver stage” (Fig. 2E) (10)(11)(14). Importantly, the predicted targets contained broadly genes expressed in the sporozoite, not only genes belonging to the groups “sporogony” and “egress oocyst and salivary-gland invasion”, which may have expression peaks in the oocyst/oocyst sporozoite used for ChIP-seq, but also genes belonging to the groups “migration to the liver and hepatocyte invasion” and “development into the liver stage”, which may have expression peaks in the salivary gland sporozoite. Additionally, each group contained comprehensively genes that can belong to the group. These results suggest that AP2-Sp determines the repertoire of gene expression during the sporozoite stage and already binds to the promoters of these genes in the oocyst/oocyst sporozoite regardless of when these genes display their highest expression during the sporozoite stage.

### Predicted targets constitute a major proportion of the genes highly expressed in sporozoites

To determine the importance of AP2-Sp in transcriptional regulation during the sporozoite stage, the abundance of target gene transcripts in the sporozoite transcriptome was investigated. RNA-seq analysis was performed in the oocyst/oocyst sporozoite (infected midguts at 14 days post-infection) and salivary gland sporozoites (sporozoites at 24 days post-infection), and the *P. berghei* genes were ordered by expression levels based on RPKM (Reads Per Kilobase of exon per Million mapped reads) values (Table 4A and Table S4B). In the oocyst/oocyst sporozoite stage, 69 genes were targets among the top 100 genes (Fig. 3A, left). These genes included two that were revealed to be putative targets by detailed manual examinations; one gene harbored peaks more than 1,200 bp upstream with accompanying transcripts downstream of the peak, while the other was not predicted to be a target due to alternative transcription or incorrect annotation. These results suggest that targets constitute a major proportion of the highly expressed genes during this stage. Among 31 non-target genes, 24 were genes related to protein synthesis including 19 genes encoding ribosomal proteins. In the salivary gland sporozoite transcriptome, among the top 100 genes, 94 were targets (Fig. 3A, right), including 13 genes that were manually predicted as targets. Eleven genes and two genes were not predicted to be targets due to the 1,200-bp criterion and presumably incorrect annotation of the genes, respectively. Three out of six non-target genes were genes related to protein synthesis including two ribosomal proteins. The high proportion of ribosomal proteins in the non-target genes in the oocyst/oocyst sporozoite transcriptome suggest that transcripts derived from oocysts before sporogony, which may abundantly produce proteins, are included in the transcriptome. This explains why the proportion of targets was higher in the salivary gland sporozoite than in the oocyst/oocyst sporozoite. Fig. 3B shows the relationship between the proportions of target genes and gene expression levels (RPKM values). The proportion of targets increased with RPKM values. These results suggest that the targets of AP2-Sp constitute a major proportion of the highly transcribed genes both in the oocyst/oocyst sporozoite and in the salivary gland sporozoite.

**Figure 3.**
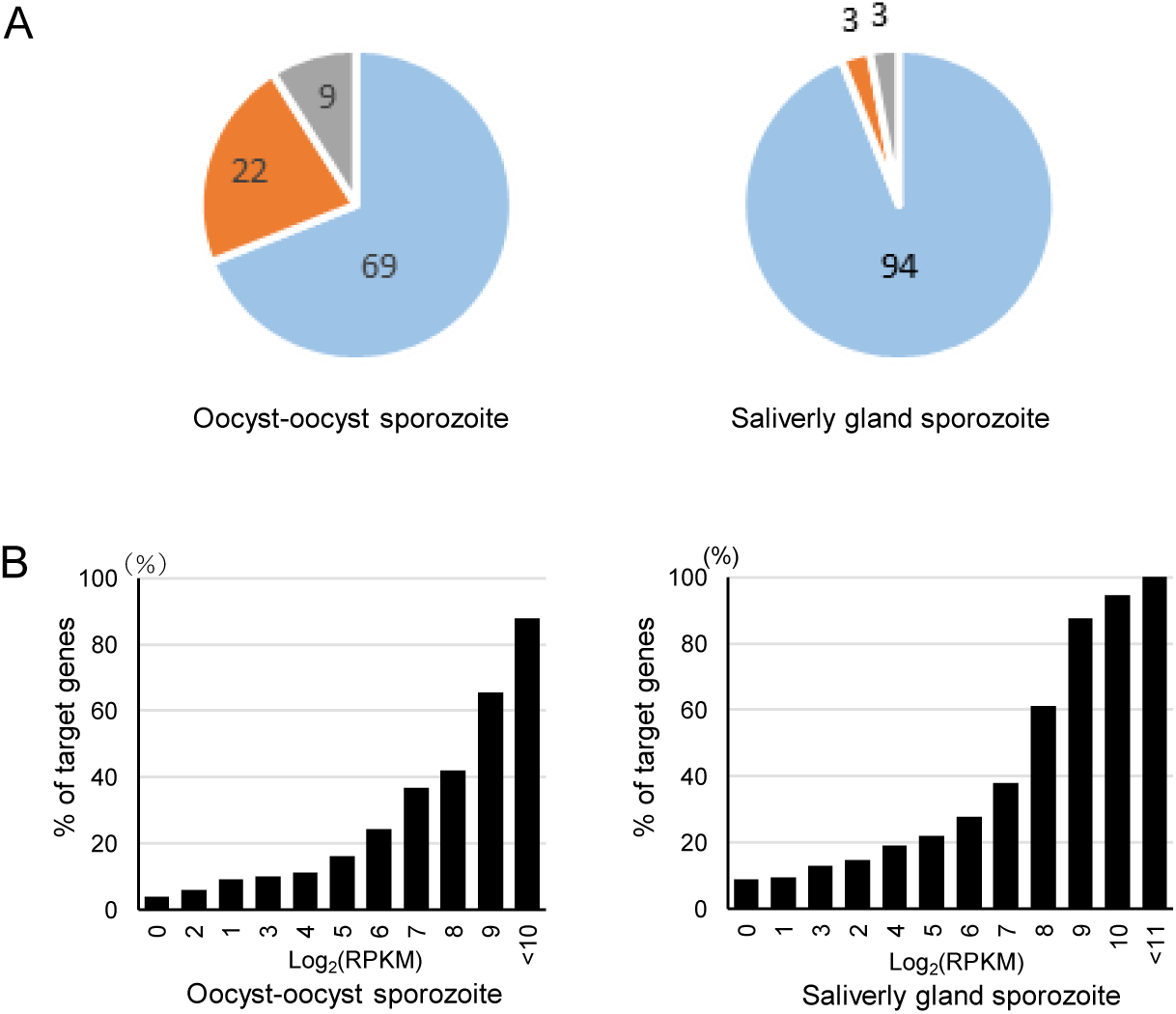
Targetome of AP2-Sp in sporozoite transcriptome. A. *Plasmodium berghei* genes excluding putative subtelomeric genes (4,654 genes) were ordered according to the RPKM values obtained by RNA-seq analyses in the oocyst/oocyst sporozoite (14 days after infective blood meal) and in the salivary gland sporozoite (21 days after infective blood meal). The proportion of target genes in the top 100 highly expressed genes is shown as a pie graph for each transcriptome. Non-target genes are divided into two groups: genes related to protein synthesis and other genes. The number of target genes in each group is shown on a pie graph. B. *P. berghei* genes were grouped according to RPKM values (in log_2_ scale), and the percentage of target genes in each group is plotted in a bar graph.

### AP2-Sp activates its transcription through a transcriptional positive feedback loop

Master TFs in the development of multicellular organisms are thought to stabilize their own expression by transcriptional auto-activation mechanisms such as transcriptional positive feedback loops, which further stabilize expression of the target genes, contributing to establishment of the cell type(15). A similar mechanism may be employed in gametocytogenesis of plasmodium parasites because AP2-G (a master regulator of gametocytogenesis) harbors binding sites of itself in its own upstream regulatory region (16). In the *P. berghei AP2-G* gene, ChIP-seq peaks for AP2-G extend across a 2-kbp upstream region beyond the ordinary promoter size (1,200 bp) that we assume in this parasite (17). To investigate whether this is true for AP2-Sp as well, we investigated AP2-Sp ChIP-seq peaks in the upstream regulatory regions of the *AP2-Sp* gene. As shown in the graphical view in Fig. 4A, in the upstream regions of *AP2-Sp* there is a large zone harboring multiple ChIP-seq peaks extending between 1,940 bp and 2,940 bp upstream of the gene and a corresponding cluster of AP2-Sp binding motifs (23 binding motifs in the region including overlapping motifs). The RNA-seq data showed that a cluster of AP2-Sp transcripts initiates downstream of this area, indicating that this is a regulatory region for the *AP2-Sp* gene. These observations suggest that AP2-Sp utilizes a transcriptional positive feedback loop, as observed with AP2-G, while AP2-Sp itself was not predicted to be a target due to the 1,200-bp inclusion criterion.

**Figure 4.**
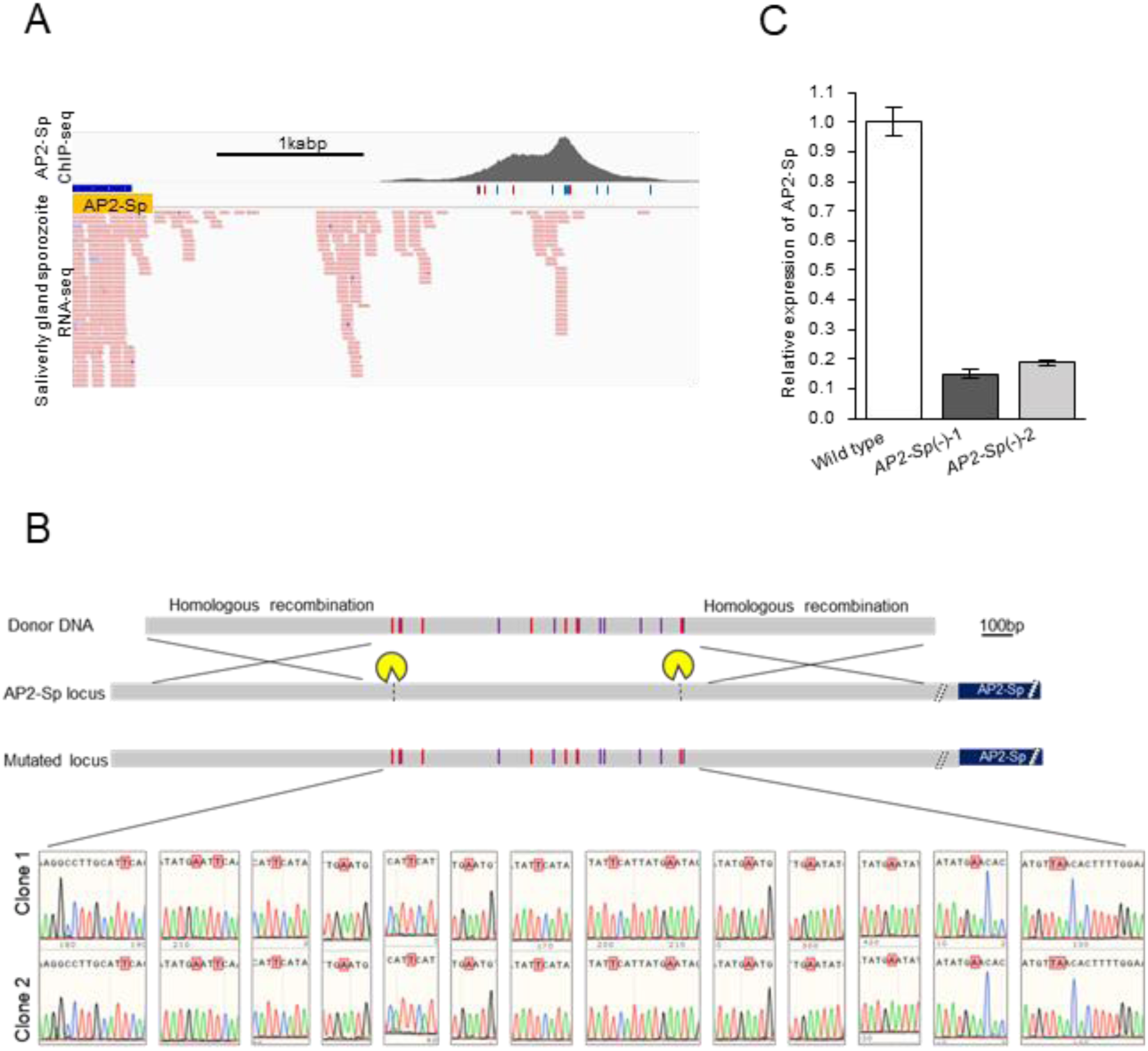
Positive transcriptional feedback is essential for maintaining AP2-Sp expression. A. Clusters of AP2-Sp peaks and corresponding putative binding motifs in the upstream region of the AP2-Sp gene. Binding motifs of AP2-Sp under the peaks are indicated by bars. Transcripts of AP2-Sp obtained by RNA-seq were mapped onto the genome parallel to ChIP-seq data. Blue and red rectangles indicate reads mapped to forward and reverse strands of the *Plasmodium berghei* genome, respectively. The image was created using Integrative Genomics Viewer (IGV) software. B. Schematic diagram of AP2-Sp_pro_mut parasite preparation. Mutant parasites were prepared from pbcas9 using the double CRISPR method. Positions targeted by gRNA are indicated by scissor characters. Vertical bars indicate mutated binding motifs of AP2-Sp (23 motifs including overlapping motifs). They were mutated by adding 16 mutations (basically, [T/C]GCA[T/C][G/A] was changed to TGAA[T/C][G/A]), which were confirmed by sequencing (lower panels). C. RT-qPCR assays of AP2-Sp transcripts were performed for pbcas9 and AP2-Sp_pro_mut parasites. Total RNA was extracted from 20 mosquito midguts at 14 days after infective blood meal. The 60S ribosomal protein L 21 mRNA was used as an internal control. Results are shown as expression of AP2-Sp relative to that in pbcas9. Data are averages of three biologically independent experiments (± SE).

We wanted to prepare parasites with all the motifs mutated to investigate if this putative positive feedback loop actually contributes to *AP2-Sp* transcription. Because the motifs were spread over a large region, it was difficult to mutate all of them simultaneously by standard genetic modification methods. Thus, the entire region was replaced with a synthetic 1,000-bp DNA fragment with mutated motifs using a double CRISPR method (Fig. 4B)(18). Two independent clones were prepared and the effects of these mutations on sporogony were assessed in mosquitoes. The oocyst number formed on the mosquito midgut wall was normal in these parasites, but the number of oocyst sporozoites obtained from them decreased significantly compared to the original parasite, pbcas9 (Table 1). The *AP2-Sp* transcripts extracted from infected midguts were five to seven times lower in both clones than in pbcas9 (Fig. 4C). These results demonstrate that a positive transcriptional feedback is important for maintaining AP2-Sp expression at a level sufficient for the normal progression of sporogony.

**Table 1.**
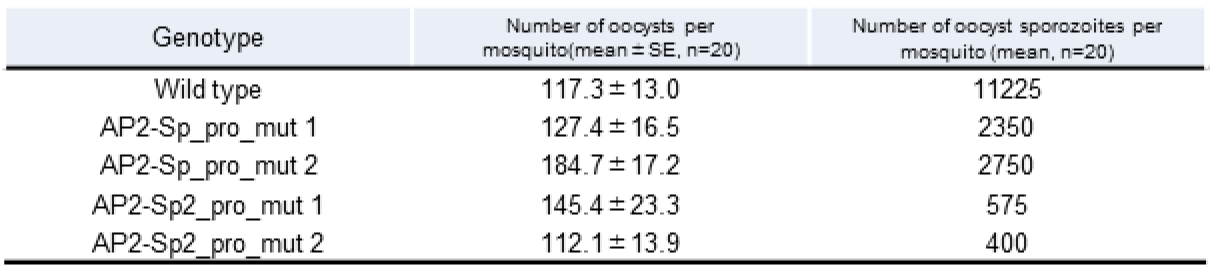
Results of mosquito transmission experiments in AP2-Sp_pro_mut and AP2-Sp2_pro_mut parasites. For each transcription factor, two biologically independent mutant parasites, clone 1 and clone 2, were prepared from pbcas9 and used for experiments. The numbers of oocysts and oocyst sporozoites were compared between pbcas9 and mutant parasites 14 days after infective blood meal. SE, standard error

### Transcriptional cascade in sporozoites is triggered by AP2-Sp

AP2 family TFs sometimes have large upstream regulatory regions, as observed in AP2-Sp and AP2-G, and are not predicted to be targets by the 1,200-bp criterion. In sporozoites, four TFs have been reported to be transcribed: the AP2 family proteins, AP2-Sp2, AP2-Sp3, and AP2-L, and the unclassifiable TF SLARP (3)(19)(12). Except for AP2-Sp2 (Fig. 4A), these TFs were not predicted to be targets of AP2-Sp. However, manual investigation of the upstream regions of these genes revealed that all harbored AP2-Sp peaks, which accompanied their downstream transcripts (Fig. 4B-4D). These results suggest that AP2-Sp regulates these genes directly.

Next, we prepared parasites in which the AP2-Sp binding motifs were mutated in the upstream region of the *AP2-Sp2* gene and examined whether *AP2-Sp2* was directly activated by AP2-Sp. The AP2-Sp2 gene harbors multiple AP2-Sp peaks in its upstream region, with nine binding motifs under the peaks (Fig. 4A). We intended to mutate all of these motifs using a double CRISPR system by targeting those nearest to each end of the peak region, but protospacer adjacent motif (PAM) sequences were not present around the motif at the 3′ end of the peak region. Thus, the third motif from this end was targeted by gRNA (Fig. 5E). Two independent clones were obtained. Two motifs near the 3′-end of the peak region of one of the clones were not mutated probably because of crossover homologous recombination between the second and third motifs. Despite these differences, the phenotypes were essentially the same in both clones. Oocysts formed as in wild type parasites, but the number of sporozoites collected from the midguts was approximately five times lower in the mutants than in the wild type (Table 1). RT-qPCR analysis showed that transcripts of *AP2-Sp2* decreased significantly (Fig. 5F). These results demonstrate that AP2-Sp activates *AP2-Sp2* directly through the seven binding sites and that this direct regulation is essential for the normal progression of sporogony.

**Fig. 5.**
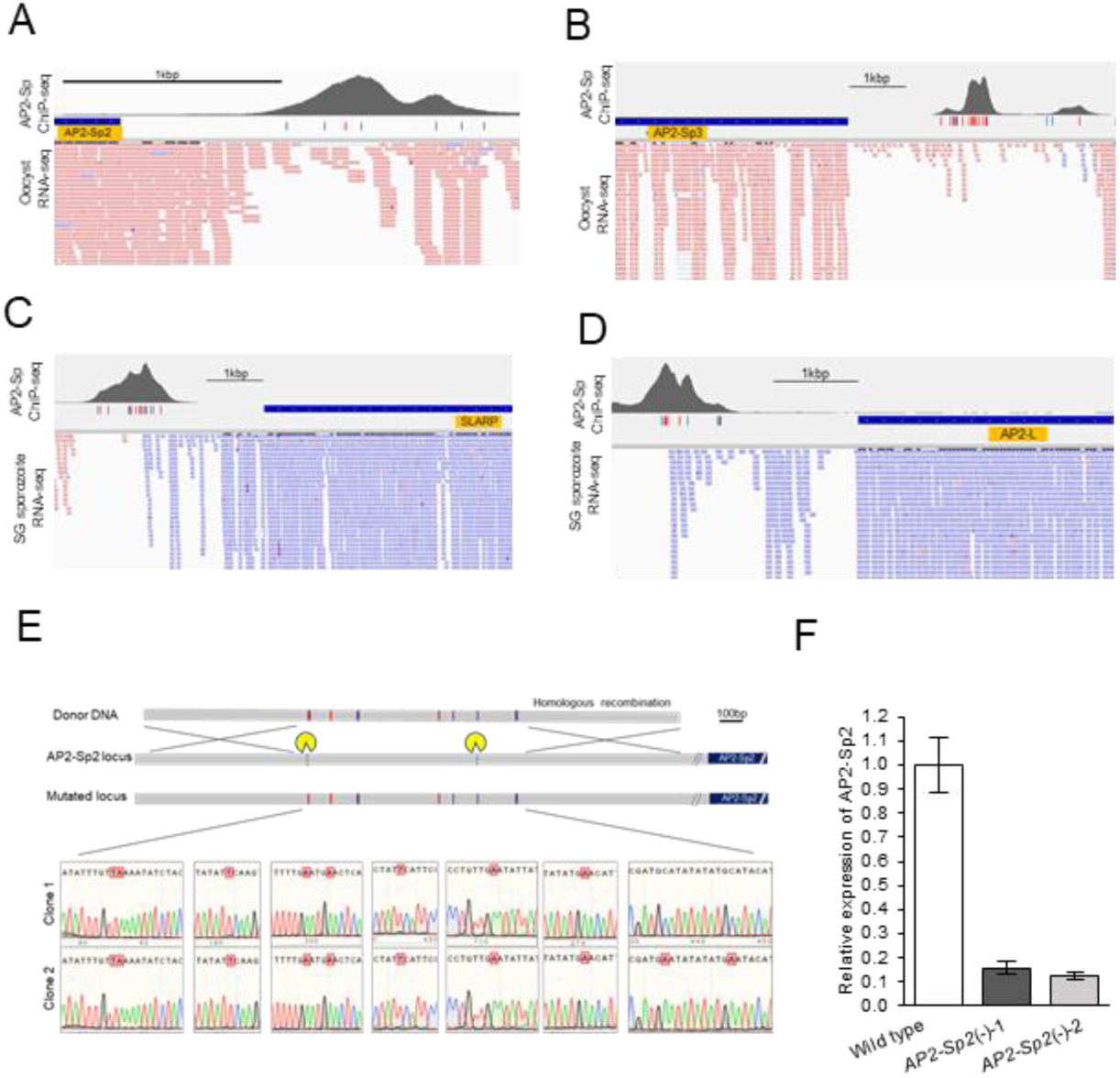
TFs specific to the sporozoite stage are targets of AP2-Sp. A. Clusters of AP2-Sp peaks and putative binding motifs of AP2-Sp under the peaks in the upstream region of the AP2-Sp2 gene. Binding motifs of AP2-Sp under the peaks are indicated by bars. Transcripts of AP2-Sp obtained by RNA-seq were mapped onto the genome parallel to ChIP-seq data. Blue and red rectangles indicate reads mapped to forward and reverse strands of the *P. berghei* genome, respectively. The image was created using Integrative Genomics Viewer (IGV) software. B. Clusters of AP2-Sp peaks and putative binding motifs of AP2-Sp in the upstream region of the AP2-Sp3 gene. C. Clusters of AP2-Sp peaks and putative binding motifs of AP2-Sp in the upstream region of the SLARP gene. D. Clusters of AP2-Sp peaks and putative binding motifs of AP2-Sp in the upstream region of the AP2-L gene E. Schematic diagram of the preparation of AP2-Sp2_pro_mut parasites. Mutant parasites were prepared from pbcas9 using the double CRISPR method. Positions targeted by gRNA are indicated by scissor characters. Vertical bars indicate mutated binding motifs of AP2-Sp. Basically, [T/C]GCA[T/C][G/A] was changed to TGAA[T/C][G/A]. Mutations were confirmed by sequencing (lower panels). Two clones were prepared independently, in which seven (clone 1) and nine (clone 2) motifs (not including overlapped ones) were found to be mutated. F. RT-qPCR assays of AP2-Sp2 transcripts. Experiments were performed between pbcas9 and AP2-Sp2_pro_mut parasites at 14 days after infective blood meal using 20 mosquito midguts as starting material. The 60S ribosomal protein L 21 mRNA was used as an internal control. Results are shown as expression of AP2-Sp2 relative to that in pbcas9. Data are averages of three biologically independent experiments (± SE).

## Discussion

The mechanisms of gene regulation in the sporozoite stage have been elusive. AP2-Sp is a candidate master regulator of this stage; it is essential for sporogony, it is expressed throughout the sporozoite stage, and many genes important for sporozoite functions harbor its binding motif in their upstream regulatory regions. However, genome-wide studies of its target genes have not been performed due to difficulties in preparing sufficient numbers of parasites. To solve this problem, we explored ways to perform ChIP-seq on this stage and finally determined that high quality data could be obtained using infected mosquito midguts as the starting material. The data revealed that this TF is a master TF of this stage with two prominent features. First, the target genes of this TF constitute a major proportion of the genes that are highly expressed during this stage and comprehensively represent genes known to be important for stage-specific functions. This style of gene regulation, which we called “direct and comprehensive regulation” in our previous paper on gene regulation in the ookinete (another motile stage)(6), suggests that AP2-Sp binding determines the gene expression repertoire in this stage. Second, AP2-Sp stabilizes its expression by a positive transcriptional feedback loop and controls other sporozoite-specific TFs, i.e., it is at the pinnacle of transcriptional regulation during this stage. The mechanism of a positive transcriptional feedback loop may be important for complete stage conversion by increasing AP2-Sp transcripts rapidly and stabilizing new gene expression repertoires that are constituted from its target genes. These identified features of this TF imply simple mechanisms of stage-specific gene regulation and suggest that a complete picture of gene regulation during this stage can be obtained by studying transcriptional regulation starting with AP2-Sp.

One question to be answered in the next step is how different profiles of gene expression are generated among the AP2-Sp target genes. Sporozoites target different organs in vertebrate and mosquito hosts. Consequently, the expression profiles of many genes change before and after salivary gland infection. For example, the expression of *UIS4*, which is required for liver-stage development, is low before salivary gland infection but upregulated greatly after salivary gland infection(20). In contrast, the expression of *serine repeat antigen 5*, which is required for oocyst rupture, is high in the oocyst sporozoite but decreases in the salivary gland sporozoite(21). If AP2-Sp binds to the promoters of these genes irrespective of their expression profiles, how can the different gene expression profiles observed in this stage occur? Do additional regulatory mechanisms exist?

At present, the most likely answer to this remaining question(s) is a scenario where TFs downstream of AP2-Sp play these roles, moderating the basal transcriptional activity of the promoter controlled by AP2-Sp and acting as repressors or enhancers to switch the expression pattern of a group of target genes. Indeed, considering that parasites show different phenotypes by disruption of AP2-Sp2, AP2-Sp3, and SLARP at this stage(3)(12), there is a possibility that these factors regulate different groups of genes, producing different profiles of gene expression among target genes of AP2-Sp. The present ChIP-seq method should help us to examine this assumptions, at least in AP2-SP2 and AP2-Sp3, which could be expressed in oocysts/oocyst sporozoites, by determining the target genes of these TFs and investigating if binding of these TFs explain their expression profiles. On the other hand further strategies to analyze genome binding may be required to study on gene regulation by SLARP, which is mainly expressed in the salivary gland sporozoite where lower numbers of parasites are obtained.

As discussed above, the cascade induction of TFs such as AP2-Sp2, AP2-Sp3 and SLARP could play an important role in determining gene expression patterns unique to the sporozoite stage. In contrast, the induction of AP2-L is not responsible for transcriptional regulation in the sporozoite stage but plays a role in initiating prompt conversion to the liver stage (19). This role seems to be analogous to that observed in gametocyte master TF AP2-G. AP2-G induces hundreds of genes and establishes early gametocytes, but at the same time, activates TFs that act in subsequent female gametocytes, which makes it possible to sustain sexual development that continues after expression of AP2-G is over (17). Presumably, the master TFs of this parasite play a role in triggering the conversion to subsequent stages and a series of cascade events of master TFs drives the life cycle forward. Taken together, these observations suggest two roles of a master TF in the lifecycle: create the basic gene expression pattern of a new stage and maintain it after stage conversion, and prepare the next stage by activating its master TFs.

In conclusion, this study demonstrates the role of AP2-Sp as a master regulator in the sporozoite stage and, to the best of our knowledge, is the first to reveal an overall picture of transcriptional regulation in this stage.

## Materials and Methods

### Ethics statement

This study was performed according to the recommendations in the Guide for the Care and Use of Laboratory Animals of the National Institutes of Health in order to minimize animal suffering. These experiments were approved by the Animal Research Ethics Committee of Mie University (permit number 23–29).

### Parasite preparations

The ANKA strain of *P. berghei* was used in all experiments. For infection of mosquitoes, infected BALB/c mice were subjected to *A. stephensi* mosquitoes. Fully engorged mosquitoes were selected and maintained at 20 °C in a fumed environment.

### ChIP-Seq

The mosquito midgut was excised, fixed immediately in 1% paraformaldehyde, and stored in this solution until the dissections of all mosquitoes were completed. ChIP-seq was performed using the same procedure as reported previously. Briefly, the fixed midguts were sonicated with a Bioruptor (Tosho Denki, Yokohama, Japan) in 0.25 ml of lysis buffer. After centrifugation at 4 °C, the supernatant was subjected to ChIP using anti-GFP antibodies, and the harvested DNA fragments were subjected to sequencing. Input DNAs were obtained from the chromatin by the same procedure but without IP. Sequencing was performed using the SOLiD 5500 System (Life Technologies) in experiment 1 and Illumina NextSeq System in experiment 2. The data have been deposited in the Gene Expression Omnibus (GEO) database under accession number GSE188985.

### Analysis of ChIP-Seq data

The sequence data were mapped onto the *P. berghei* genomic sequence (PlasmoDB, version 3) and the *A. stephensi* genome (IndV3) (22) using Bowtie software under default settings. The mapping data of *P. berghei* were further analyzed using the MACS2 peak-calling algorithm (false discovery rate (FDR)<0.01). To identify common peaks between two experiments, peaks summits within 150 bp were selected using an in-house script. To estimate the binding sequences of AP2-Sp, six-base sequences concentrated within 100 bp from the predicted summits were investigated by an in-house script using Fisher’s exact test as described previously(6). Genes were determined to be targets of AP2-Sp when they were located within 1,200 bp downstream of the predicted summits of peaks. When the upstream region was less than 1,200 bp, the whole intergenic region was used instead for predictions.

### RNA-seq

For the RNA-seq of theoocyst/oocyst sporozoite, total RNA was extracted from the midguts dissected from ten mosquitoes 14 d after an infective blood meal. The dissected midguts were soaked in RNAlater solution until all midguts had been collected. Ten midguts were used for each extraction, and three independent experiments were performed. For the RNA-seq of salivary gland sporozoites, total RNA was extracted from salivary gland sporozoites 21 d after an infective blood meal. One hundred mosquitoes were used for each extraction (usually approximately one million sporozoites were obtained), and three independent experiments were performed. RNA was extracted using Isogen II (Nippon Gene) according to the manufacturer’s protocol. The dissected midguts were lysed in the lysis solution supplied with the kit using the TissueLyser system (Qiagen). The library for RNA-seq was prepared using a HyperPrep kit (KAPA) and then used for Illumina sequencing. Reads were mapped on the *P. berghei* genome with HISAT2 software. The HISAT2 parameters were set to default except for “--max-intron”, which was set to 1,000. The mapping data were analyzed by featureCounts software to calculate RPKM values. Average RPKM values obtained by three independent experiments were used to quantify gene expression.

### Preparation of mutant parasites by CRISPR/Cas9 system

Parasites with mutated AP2-Sp binding motifs were prepared with the CRISPR/Cas9 system using *P. berghei* parasites constitutively expressing *Cas9* (pbcas9), as reported previously(18). A DNA fragment corresponding to the upstream region of the gene was synthesized with all motifs mutated (two point mutations per motif). Sequences for homologous recombination were added to both sides of the fragment by overlapping PCR and used as donor DNA. Two guide RNA (gRNA) sequences were designed, corresponding to the motifs on opposite sides of the synthesized fragment. The gRNA target sequences were subcloned into a plasmid for double CRISPR gRNA expression. Mature schizonts were transfected with donor DNA and the gRNA plasmid, and mutants were selected with pyrimethamine for 1.5 d, beginning 30 hours after transfection. Parasite clones were obtained by limiting dilution, and correct exchange of the original locus with the donor DNA fragment by double-crossover homologous recombination was verified by sequencing. The primers used in this experiment are listed in Table S5.

### Reverse transcription-quantitative PCR (RT-qPCR) assay

Twenty midguts were dissected 14 d after an infective blood meal. Total RNA was extracted from the excised midguts using the same procedures as described above. cDNA was synthesized using a PrimeScript RT Reagent Kit with gDNA Eraser (Takara). The RT-qPCR assay was performed using the TB Green Fast qPCR Mix (Takara). The ribosomal protein L21 gene (*RPL21*, PBANKA_1018600) was used as an internal control. The primers used in this assay are listed in Table S5.

## Acknowledgments

This work was supported by JSPS KAKENHI Grant Number 17H01542 to MY.

## Supporting information captions

### Supplementary tables

Table S1 (separate file). Summary of experiment 1.

Table S1A. AP2-Sp peaks identified in experiment 1. Peaks were called by MACS2 software with q-value < 0.01, and further selected by values of fold enrichment (>3).

Table S1B. Six-base sequences enriched around the summits of AP2-Sp peaks in Table S1A. AP2-Sp peaks identified in experiment 1. Six-base sequences enriched around the summits of peaks (within 100-bp from summits) identified in experiment 1 (Table S1A) were calculated. They are arranged beginning with the sequence with the smallest p-value.

Table S1C. Targets of AP2-Sp predicted from AP2-Sp peaks identified in experiment 1.

Table S2 (separate file). Summary of experiment 2.

Table S2A. AP2-Sp peaks identified in experiment 2. Peaks were called by MACS2 software with q-value < 0.01, and further selected by values of fold enrichment (>3).

Table S2B. Six-base sequences enriched around the summits of AP2-Sp peaks in Table S2A. Six-base sequences enriched around the summits of peaks (within 100-bp from summits) identified in experiment 2 (Table S2A) were calculated. They are arranged beginning with the sequence with the smallest p-value.

Table S2C. Targets of AP2-Sp predicted from AP2-Sp peaks identified in experiment 2.

Table S3 (separate file). Common targets and their classification.

Table S3A. Common targets between those predicted by experiment 1 and those predicted by experiment 2

Table S3B. Classification of AP2-Sp target genes. Target genes of AP2-Sp common between those predicted by experiment 1 and those predicted by experiment 2 were classified according to the functional annotation in PlasmoDB.

Table S4 (separate file). Target genes in the top 500 genes highly expressed in the sporozoite. Table S4A. Target genes in the top 500 genes highly expressed in the oocyst/oocyst sporozoite. Plasmodium berghei genes were ordered according to their expression levels (RPKM values) based on RNA-seq data in the oocyst/oocyst sporozoite.

Table S4B. Target genes in the top 500 genes highly expressed in the salivary gland sporozoite. Plasmodium berghei genes were ordered according to their expression levels (RPKM values) based on RNA-seq data in the salivary gland sporozoite.

Table S5 (separate file). Primers used in this study.

